# Collaborative Cross Mouse Populations as a Resource for the Study of Epilepsy

**DOI:** 10.1101/690917

**Authors:** Bin Gu, John R. Shorter, Lucy H. Williams, Timothy A. Bell, Pablo Hock, Katherine A. Dalton, Yiyun Pan, Darla R. Miller, Ginger D. Shaw, Brian C. Cooley, Benjamin D. Philpot, Fernando Pardo-Manuel de Villena

## Abstract

Epilepsy is a neurological disorder with complex etiologies and genetic architecture. Animal models have a critical role in understanding the pathophysiology of epilepsy. Here we studied epilepsy utilizing a genetic reference population of Collaborative Cross (CC) mice with publicly available whole genome sequences. We measured multiple epilepsy traits in 35 CC strains, and we identified novel animal models that exhibit extreme outcomes in seizure susceptibility, seizure propagation, epileptogenesis, and sudden unexpected death in epilepsy. We performed QTL mapping in an F2 population and identified seven novel and one previously identified loci associated with seizure sensitivity. We combined whole genome sequence and hippocampal gene expression to pinpoint biologically plausible candidate genes and candidate variants associated with seizure sensitivity. These resources provide a powerful toolbox for studying complex features of seizures and for identifying genes associated with particular seizure outcomes, and hence will facilitate the development of new therapeutic targets for epilepsy.

---

Epilepsies are a clinically heterogeneous group of neurological disorders affecting ∼1% of the worldwide population^1^. There are no treatments to prevent epilepsy, and roughly 30% of epilepsies are intractable to current antiepileptic medications^2^. Uncontrolled seizures also increase the risk of sudden unexpected death in epilepsy (SUDEP), a poorly understood fatal complication of epilepsy^3^. New transformative treatments could be developed by understanding the genetic associations conferring risk or protective effects. In addition to rapid expansion of epilepsy-associated gene list discovered mainly with monogenic causes of seizures, human GWAS studies of generalized epilepsy identified more than a dozen novel genome-wide significant loci and biological plausible candidate genes^4-6^. However, human GWAS approaches have had limited success in identifying risk loci associated with certain forms of epilepsy (e.g. partial epilepsy) and specific seizure outcomes (e.g. SUDEP), partially due to complex disease etiologies and underpowered sample sizes^7,8.^ Animal studies remain essential for understanding the mechanisms of epilepsy and for identifying new therapeutic targets. In particular, animal models offer phenotypic repeatability and a means to study a trait in a controlled environment, as well as providing a level of experimental testing and validation that is not ethical or possible in human subjects, such as in the case of SUDEP. Most existing animal models of complex diseases such as epilepsy are limited because they do not mirror the genetic diversity found in the human population. In order to identify novel genetic targets, a model research population with a high level of genetic variation is needed.

The Collaborative Cross (CC) population is a genetically diverse recombinant inbred panel derived from eight fully inbred strains that has ∼42 million segregating genetic variants^9^. In addition, whole genome sequences have been established for each individual CC strain, and other genomic tools have been developed to enable the identification of genetic variants from mapping studies^9,10^. The CC has been used to model complex traits such as behavior^11-14^, cancer^15^, and infectious disease susceptibility^16-23^. In order for us to take advantage of the immense genetic diversity available in the CC to study epilepsy, we needed a seizure induction protocol amenable to high throughput analysis. Accordingly, we studied the CC using flurothyl seizure-induction to characterize multiple epileptic outcomes including seizure susceptibility, seizure propagation, seizure development (*i.e*. epileptogenesis), and SUDEP. We outlined a general approach to use the CC and its associated genetic resources to identify quantitative trait loci (QTL) and candidate genes. We demonstrate a highly reliable and repeatable seizure induction paradigm, as well as the implementation and design of a genetic mapping population. We also show the power of our approach by using the CC and transcriptomic sequence resources to identify several QTL (both novel and previously identified), candidate genes, and candidate variants associated with seizure sensitivity. These resources provide a powerful toolbox for identifying new genes, and hence new therapeutic targets, linked to seizure susceptibility, seizure propagation, epileptogenesis, and SUDEP.

## RESULTS

### Extreme seizure responses in CC strains

Previous studies have reported that classical inbred strains of mice vary in their naïve responses to seizures^24-26^. These studies have led to the general classification of seizure resistant (e.g. C57BL/6J, B6J) and seizure susceptible (e.g. DBA/2J) mouse strains^24-26^. Here, we used the flurothyl model to evaluate seizure-related phenotypes in a sampling of 35 CC inbred strains in addition to canonical seizure resistant (i.e. B6J) and susceptible (i.e. DBA/2J) inbred mice (Fig. 1a). We observed wide variation (∼2-fold difference) for the latency to the onset of myoclonic seizure (Myoclonic seizure threshold, MST; CC037: 320.7±16.1 sec vs. CC027: 164.6±8.2 sec) and generalized seizure (Generalized seizure threshold, GST; CC058: 472.7±23.9 sec vs. CC031: 191.2±12.9 sec). This variation extends beyond the canonically resistant and particularly beyond susceptible strains (Fig. 1b,c). We also estimated narrow sense heritability (h^2^) and observed that both traits exhibited high heritability (MST = 0.63, GST = 0.71), indicating a strong additive genetic component for flurothyl induced seizure sensitivity in this population.

**Figure 1.**
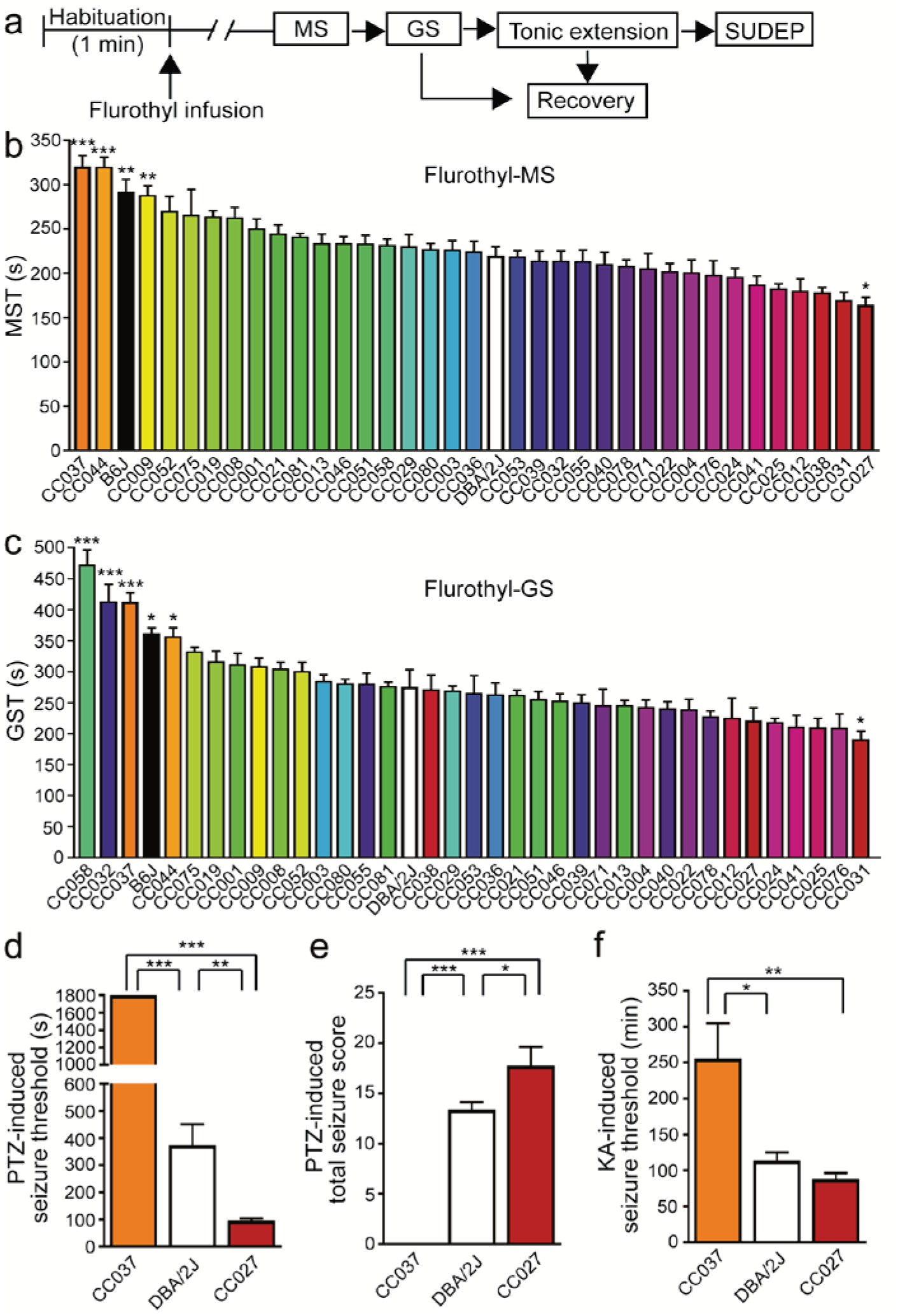
Seizure thresholds and severity highly depend upon genetic background. **a**, Schematic of flurothyl-induced seizures. MS = myoclonic seizure; GS = generalized seizure; SUDEP = sudden unexpected death in epilepsy. **b**, MS threshold (MST) and **c**, GS threshold (GST) of 35 CC strains as well as B6J and DBA/2J. The order of strains was ranked from the most resistant (highest threshold) to the most susceptible (lowest threshold), n=3–10. Data are presented as mean ± SEM and analyzed using one-way ANOVA with *post hoc* Dunnett’s multiple comparisons test. *p<0.05, **p<0.01 and ***p<0.001 compared to DBA/2J. **d**, Seizure threshold and **e**, sum of maximum behavioral seizure score after PTZ injection (40 mg/kg, i.p.), n=5. **f**, Latency to onset of generalized seizures induced by repeated low dose of KA (5 mg/kg, i.p. every 30 min), n=4-5. Data are presented as mean ± SEM and analyzed using one-way ANOVA with *post hoc* Tukey’s multiple comparisons test. *p<0.05, **p<0.01 and ***p<0.001.

To test whether the flurothyl seizure susceptibility phenotypes were generalizable to other seizure models, we tested CC037 (highly resistant), DBA/2J (intermediate), and CC027 (highly susceptible) strains in two additional seizure induction paradigms — intraperitoneal injection of pentylenetetrazol (PTZ) and kainic acid (KA), as these paradigms trigger seizures via different routes and/or mechanisms^27^. We found that the direction of effect and the significant difference of seizure responses is consistent between flurothyl- (MST, *p*-value<0.001; GST, *p*-value<0.001), PTZ- (threshold, *p*-value<0.001; seizure score, *p*-value<0.001), and KA-induced (*p*-value<0.01, one-way ANOVA) seizure models (Fig. 1d–f), demonstrating that the seizure responses of CCs measured by flurothyl can be generalizable regardless of induction paradigm.

Previous research reported evidence of a significant correlation between MST and GST across multiple standard laboratory strains in their first exposure to flurothyl, suggesting an interaction between these two phenotypes^28^. Consistently, we observed a strong correlation of initial MST and GST (R^2^=0.825, *p*-value<0.001) across most CC strains as well as B6J and DBA/2J strains (Fig. 2a). However, two CC strains, CC058 and CC032, exhibited a significant prolonged MS-GS interval compared to other strains (*p*-value<0.001) (Fig. 2b). This suggests that there are shared biological processes for sensitivity for MST and GST, but these factors can be decoupled as shown in CC058 and CC032. This also raises the possibility that target genes, and thus druggable targets, can be identified that halt seizure propagation and provide protective effects to having a generalized seizure.

**Figure 2.**
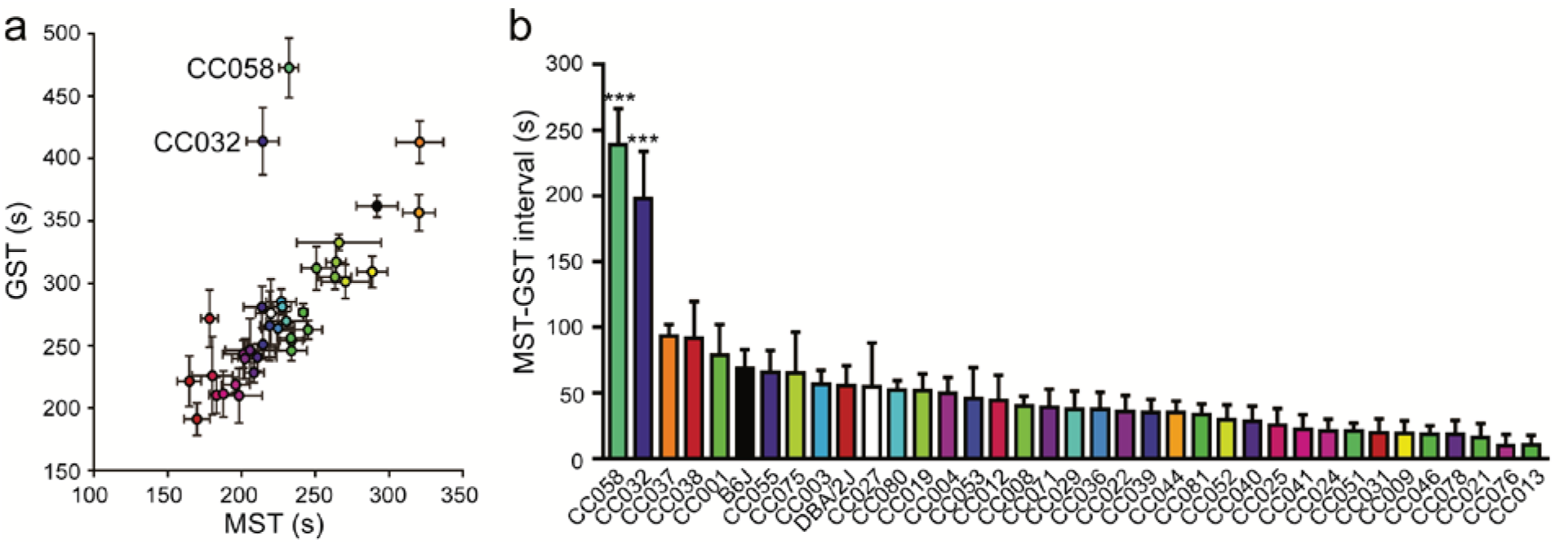
Correlation of myoclonic and generalized seizure thresholds and identification of seizure propagation resistant CC strains. **a**, Correlation of MST and GST (R^2^=0.824, p<0.001 without CC032 and CC058). **b**, MST-GST interval of 35 CC strains as well as B6J and DBA/2J, n=3-10. Data are presented as mean ± SEM and analyzed using one-way ANOVA with *post hoc* Tukey’s multiple comparisons test. ***p<0.001 compared to other strains.

### Variable kindling kinetics in CC strains

We measured the epileptogenic responses of the same 35 CC strains, B6J, and DBA/2J using an 8-day flurothyl kindling paradigm. We first found diverse MST and GST kindling kinetics for these strains that falls into four broad subgroups: (1) seizure threshold decreased throughout the kindling process (e.g. CC013); (2) threshold decreased and then plateaued (e.g. CC075 and CC052); (3) threshold was initially unchanged and then decreased (e.g. CC071); and (4) threshold remained similar throughout 8-days flurothyl kindling (e.g. CC051, GST kindling slope=2.335±2.390), suggesting a remarkable resistance to epileptogenesis (Fig. 3a,b and Supplemental Fig. 1). The kindling patterns were similar between the measurement of MST and GST, as kindling slopes of MST and GST are correlated (R^2^=0.325, *p*-value<0.001) across strains, except CC032, which exhibited the highest GST, but a more typical MST kindling slope (Fig. 3c).

**Figure 3.**
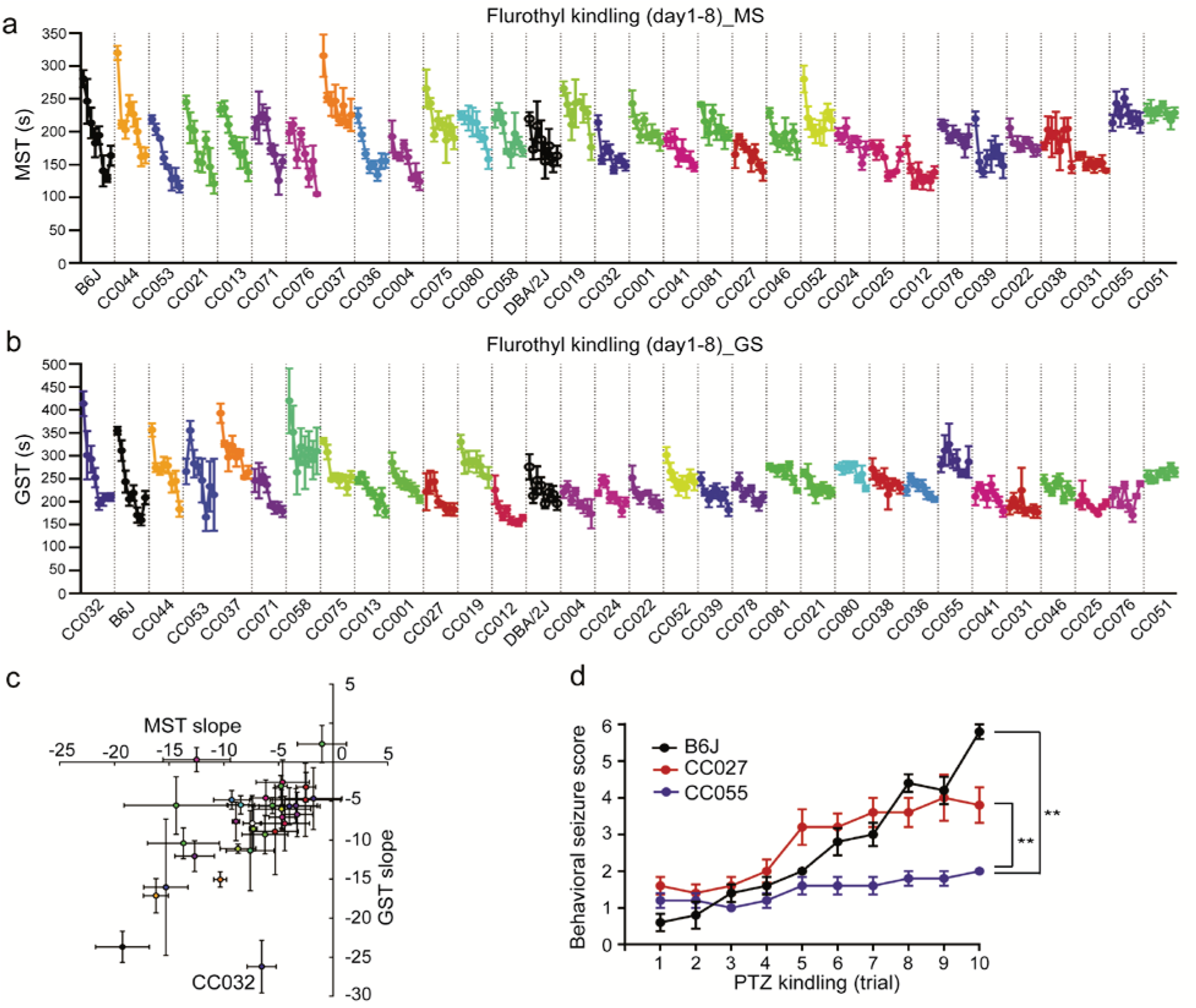
Identification of strains resistant to epileptogenesis. **a**, MST and **b**, GST during 8-day flurothyl kindling. Data of the CC strains are ordered by kindling slope. **c**, Kindling slopes of MST and GST (R^2^=0.325, p<0.001) are correlated across strains except CC032. **d**, Maximum behavioral seizure score during 30 min after each injection of PTZ (35 mg/kg, i.p.) every other day over 10-trail PTZ kindling. Data are presented as mean ± SEM and analyzed using two-way ANOVA with *post hoc* Tukey’s multiple comparisons test, n=5, **p<0.01.

To assess whether the resistance to epileptogenesis in a particular CC strain was generalizable to another induction paradigm, and based on MST kindling slope of our initial cohort (cohort 1), we challenged CC055 (lowest), B6J (highest), and CC027 (intermediate) mice to a PTZ kindling paradigm. PTZ kindling provides another measure of epileptogenesis with increases of behavioral seizure severity after repeated injection of PTZ at sub-convulsive doses^29^. Consistent with the flurothyl-kindling paradigm, behavioral seizure severity of B6J and CC027 increased over the process of PTZ kindling, whereas the seizure activity of CC055 remained sub-convulsive throughout repeated PTZ injections (*p*-value<0.01) (Fig. 3d). Overall, these results suggest that CC055 mice are similarly resistant to kindling by repeated flurothyl or PTZ exposures. Identification of an additional kindling resistant strain (i.e. CC051, lowest kindling slope) in cohort 4 offers another powerful model for resistance to epileptogenesis that could be leveraged to identify target genes.

### Induction of a single seizure significantly increases the risk of death in multiple CC strains

Flurothyl induces transient convulsions that are otherwise nonfatal to naïve standard laboratory strains^26^. During initial screening of 209 mice, 24 mice (11.5%) were unable to recover from tonic extension following flurothyl-induced seizure and succumbed to sudden death, which models SUDEP. Most (91.7%) pro-SUDEP mice were from four CC strains: CC003 (5/9), CC008 (6/10), CC009 (7/8), and CC029 (4/6). Mice from these strains that survived an initial flurothyl challenge were still susceptible to seizure-induced sudden death during subsequent flurothyl challenges (Fig. 4a). ECG recording of naïve pro-SUDEP mice as well as B6J revealed similar heart rate (Fig. 4b). However, CC009 and CC029 have a marked long QT interval (*p*-value<0.05; Fig. 4c), which is a risk factor of cardiac arrhythmia and sudden death^30^. Echocardiography also revealed CC029 has significantly reduced left ventricular fractional shortening (*p*-value<0.05; Fig. 4d), suggesting impaired myocardial contractility. We also tested whether age could be a factor for susceptibility to SUDEP. In CC009, younger mice (i.e. 2–4 months old) were significantly more susceptible to seizure-induced sudden death compared to older mice (i.e. 7–9 month old) (Fig. 4a, *p*-value<0.01, Log-rank test), similar to the enhanced susceptibility to SUDEP observed in young adults in human^31,32.^ Collectively, these results show that several mouse models for SUDEP exist in the CC, likely acting through genetically diverse mechanisms.

**Figure 4.**
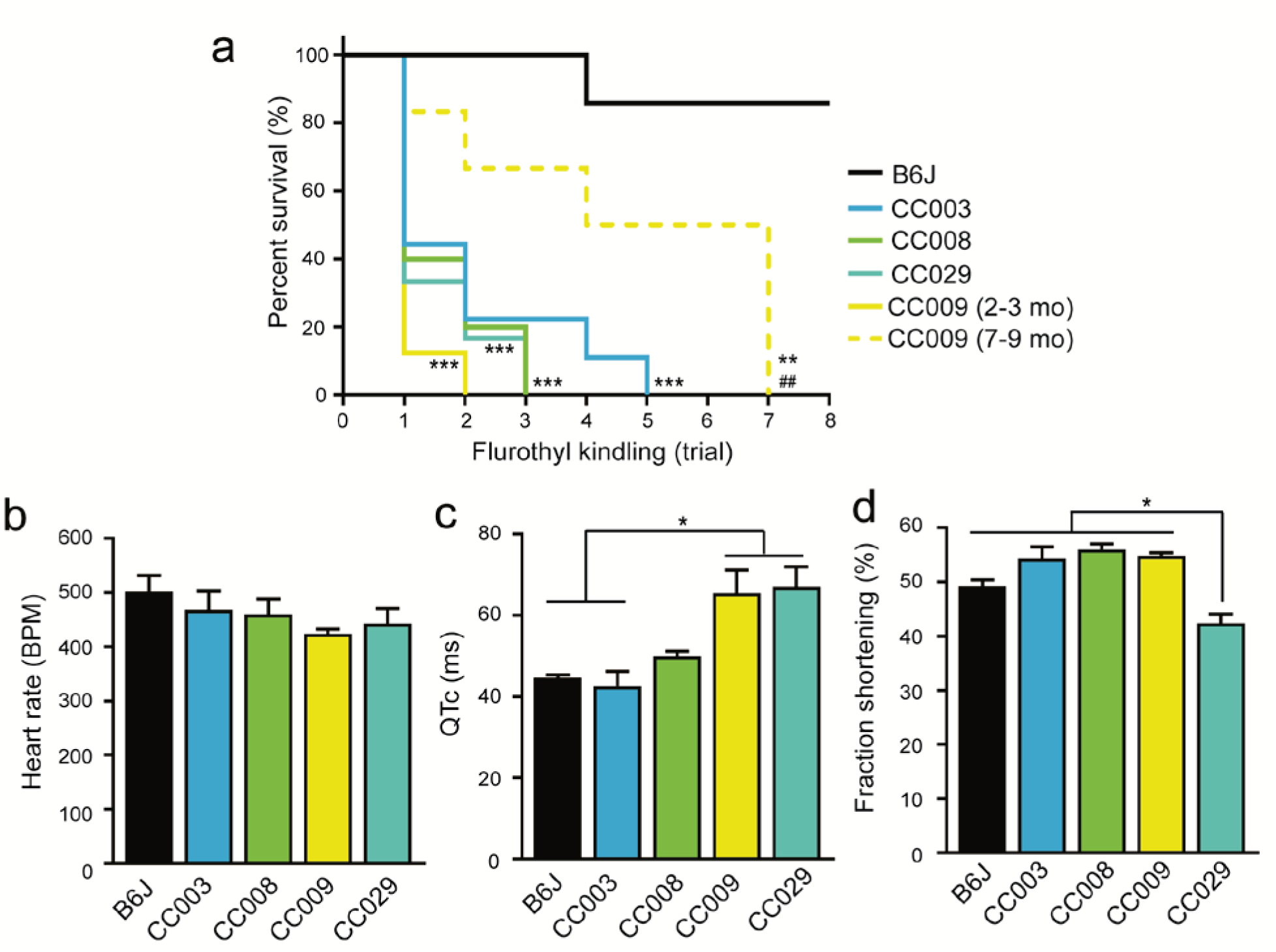
Characterization of SUDEP susceptible CC strains. **a**, Survival curve of CC003, CC008, CC009, CC029 and B6J during 8-day flurothyl kindling. Data are analyzed using Log-rank (Mantel-Cox) test, n=6–10, **p<0.01 and ***p<0.001 compared to B6J; and ^##^p<0.01 compared to young CC009 (2–3 month). **b**, Heart rate (BPM = beats per minute), **c**, corrected QT interval and **d**, left ventricular fractional shortening were measured by ECG and Echo recordings in CC003, CC008, CC009, CC029, as well as B6J mice. Data are presented as mean ± SEM and analyzed using one-way ANOVA with *post hoc* Tukey’s multiple comparisons test, n=3, *p<0.05.

### Seizure sensitivity in an F2 population

We created an F2 population to identify seizure sensitivity loci for MST and GST. CC027 and B6J strains were selected as the parental strains for an F2 mapping population for the following reasons. First, these two strains represent near phenotypic extremes for both myoclonic and generalized seizure thresholds (Figure 1). Second, B6J is the strain the mouse reference genome is based upon, and has been a canonical seizure resistant strain of choice in previous epilepsy research, allowing us to compare these results with previous work as proof of principle^24-26^. Third, CC027 is the most seizure sensitive among strains tested in cohort 1 with the lowest combined MST and GST. Fourth, CC027 exhibits similar extreme seizure sensitivities across seizure induction paradigms. Accordingly, we tested CC027 and B6J strains with both sexes represented for MST and GST and observed no significant sex differences (*p*-value=0.201 and 0.160, respectively) (Supplemental Fig. 2). We also tested F1s from CC027 and B6J mice and observed an intermediate sensitivity between the two parental strains (MST=190.7±5.1 sec; GST=265.2±9.5 sec), suggesting that additive genetic factors contribute to MST and GST (Supplemental Fig. 3).

Seizure sensitivity thresholds were measured in 297 reciprocal F2 male mice. We observed a wide range of seizure thresholds for MST (mean=176.1, SD=38.8, range: 59–278 sec) and GST (mean=273.2, SD=57.3, range: 157–540 sec) (Supplemental Fig. 3). In the F2 population, we observed a modest but significant correlation between age of mice and seizure sensitivity for both MST (R^2^=0.0517, *p*-value<0.001) and GST (R^2^=0.03, *p*-value<0.01), with younger mice being more resistant to seizure. To account for this effect, age was treated as a covariate for QTL mapping. Body weight had a non-significant effect on MST (*p*-value=0.125) and GST (*p*-value=0.09). Due to a skewed right distribution for GST and outliers with a high threshold score, GST values were normalized using a log10 transformation (Supplemental Fig. 3).

### Genetic mapping of seizure sensitivity identifies 8 QTL

We performed QTL mapping to identify genomic regions associated with MST and GST sensitivity. Mice from the F2 cross as well as the parentals and F1s were genotyped on MiniMUGA with 2,440 genetic markers that segregate between CC027 and B6J. We performed QTL mapping with age as a covariate, and identified 5 significant QTL peaks (*p*-value<0.05) and 3 suggestive peaks (*p*-value<0.1) for MST (*CC myoclonic seizure susceptibility, Ccmss*) and GST (*CC generalized seizure susceptibility, Ccgss*) (Fig. 5, Table. 1). There are two overlapping QTL for both seizure traits on chromosome 5 (*Ccmss2*/*Ccgss2*) and 10 (*Ccmss3*/*Ccgss3*). Other identified QTL are specific to either MST or GST. The allelic effects of these QTL are consistent with the parental strain effects, with the exception of *Ccmss2*/*Ccgss2* on chromosome 5, which is transgressive. This means that for chromosome 5, the seizure resistant B6J strain has a seizure sensitivity allele while CC027 has a resistance allele. *Ccgss1* on chromosome 1 overlaps the *Seizure susceptibility 1* (*Szs1*) locus which was previously identified using B6J and DBA/2J mice with consistent B6J allelic effects^33-35^. As proof of principle, we also examined whether the eight QTL identified in this study were enriched for genes associated with epilepsy in humans. We first identified human orthologs of genes within each QTL and found seven out of the eight QTL harboring nine genes in total (*NKAIN3, COQ3, GABRB1, PTPRR, SLAMF1, ATP1A2, IGSF8, KCNJ10* and *RAPGEF6*) that are relevant to human epilepsies in EpilepsyGene database (Supplemental Table 1)^36^. Notably, the marker with the most significant LOD score (=6.19) for *Ccmss2* is on chromosome 5 between *Gabrg1* and *Gabra2*. This result is consistent with a recent human GWA study of epilepsies that identified *GABRA2*^*5*^, and the fact that GABA receptors are a frequent pharmacological target for seizure sensitivity^37^. Overall, the QTL mapping results demonstrate the power and utility of the CC to map both previously identified and novel genetic loci associated with two types of seizure sensitivity.

**Table 1.** QTL mapping results.

| **Trait** | **Name** | **Chr** | **Position (Mb)** | **Size (Mb)** | **CI (Mb)** | **LOD** | **P-value** | **B6J allele** | **CC027 allele** | **Dominance** |
| --- | --- | --- | --- | --- | --- | --- | --- | --- | --- | --- |
| Myoclonic | *Ccmss1* | 4 | 31.5 | 29.4 | 10.5-39.9 | 4.49 | 0.0111 | Resistant | Sensitive | Dominant |
| Myoclonic | *Ccmss2* | 5 | 70.9 | 66.2 | 66.8-133 | 6.19 | 0.0001 | Sensitive | Resistant | Additive |
| Myoclonic | *Ccmss3* | 10 | 115.8 | 76.8 | 45.2-122 | 4.54 | 0.0105 | Resistant | Sensitive | Dominant |
| Myoclonic | *Ccmss4* | 13 | 73.3 | 48.6 | 44.6-93.2 | 3.46 | 0.1046 | Resistant | Sensitive | Dominant |
| Generalized | *Ccgss1* | 1 | 173 | 14 | 163-177 | 5.08 | 0.0037 | Resistant | Sensitive | Additive |
| Generalized | *Ccgss2* | 5 | 71.9 | 70.7 | 3-73.7 | 3.45 | 0.1151 | Sensitive | Resistant | Additive |
| Generalized | *Ccgss3* | 10 | 89.6 | 77.1 | 49.9-127 | 4.69 | 0.0085 | Resistant | Sensitive | Dominant |
| Generalized | *Ccgss4* | 11 | 51.6 | 65.6 | 10.7-76.3 | 3.79 | 0.0594 | Resistant | Sensitive | Additive |

**Figure 5.**
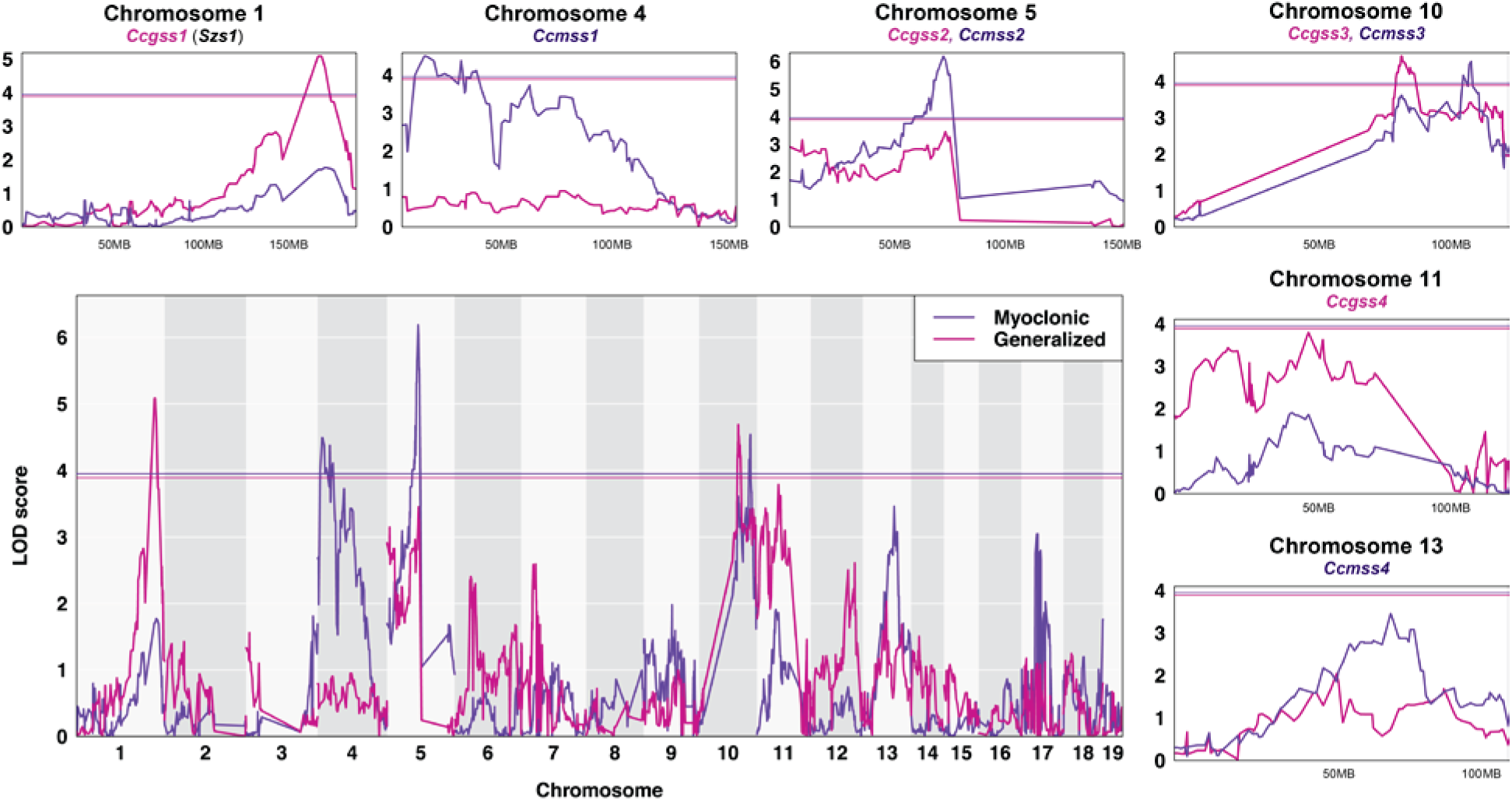
QTL mapping for myoclonic (slate-blue) and generalized (violet-red) seizure threshold. Chromosomes 1 through 19 are represented numerically on the x-axis, and the y-axis represents the LOD score. The relative width of the space allotted for each chromosome reflects the relative length of each chromosome. A magnification of each *CC myoclonic seizure susceptibility* (*Ccmss*) and *CC generalized seizure susceptibility* (*Ccgss*) locus is shown. Solid colored horizontal bar indicates the significance threshold at *p* = 0.05.

### RNA sequencing analysis

To help interrogate allelic effects of genes within QTL, we performed gene expression analysis using RNA sequencing. We selected CC027 and B6J mice to access the strain specific genotypic effects on gene expression in the hippocampus. In total, we identified 3,069 differentially expressed genes after correcting for multiple comparisons with an adjusted *p*-value <0.05. For the purposes of this analysis, we focused on gene expression within the most significant QTL, *Ccmss2*/*Ccgss2* on chromosome 5. Within a 3 Mb region surrounding the most significant marker, we identified 21 protein coding genes, of which 12 were measurably expressed in the hippocampus (Fig. 6a). Of these 12 genes, four have genome-wide differential expression (*p*-value<0.05), with *Gabra2* being the most differentially expressed (Fig. 6b). *Gabra2* in B6J has significantly decreased expression compared to CC027. In fact, *Gabra2* was one of the most differentially expressed genes in the set (*p*-value<1.0E-100).

**Figure 6.**
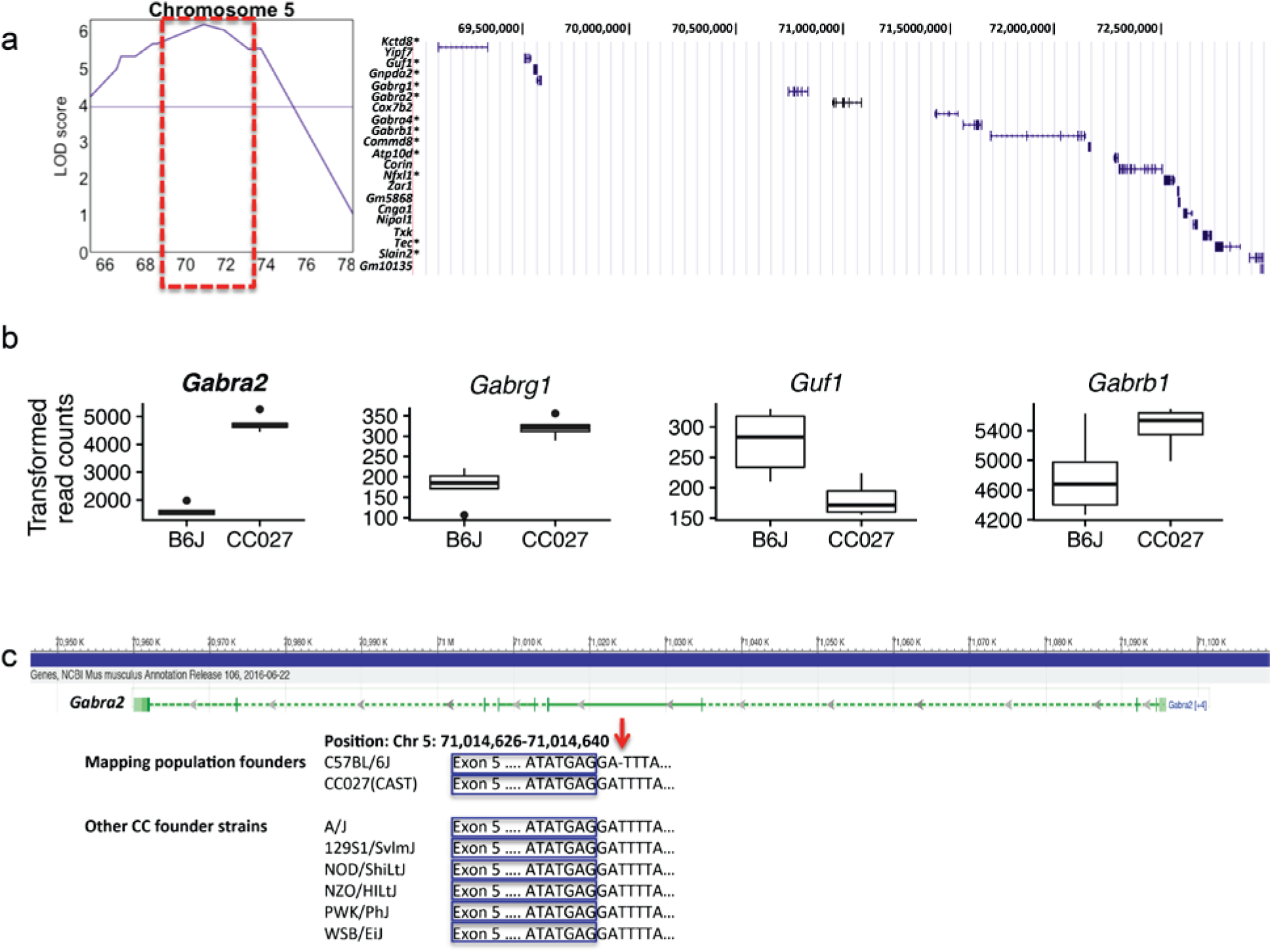
Identification of genetics variants from QTL. **a**, *Ccmss2* QTL region on chromosome 5 and list of 21 protein coding genes found within the interval. * denotes genes that were measurably expressed in the hippocampi. **b**, Genes in the QTL region with significant differential (p<0.05, t-test) expression in the hippocampus of B6J and CC027 mice. Data are presented as median with the lower and upper hinges corresponding to 25–75 percentiles. **c**, *Gabra2* with an intronic base pair deletion located next to a splice acceptor site preceding exon 5. This deletion is only found in B6J mice.

Using whole genome sequence from CC027 as well as the reference B6J sequence, we were able to identify founder haplotype and genetic variation in the exons for *Gabra2*. The origin of the region (70.95Mb–71.10Mb) in CC027 is *M. m. castanus* while in B6J this region is *M. m. domesticus*. As expected in regions with different subspecific origin, there are high levels of genetic diversity (at least 1,310 SNP variants, 181 short indels). Within the ten exons of *Gabra2*, we observed five SNPs and one indel between B6J and CC027. The indel and three of the five SNPs are UTR variants have no reported evidence for changes in gene expression. The remaining two SNPs are synonymous variants.

The most likely cause of gene expression variation in *Gabra2* comes from an intronic variant. An indel, rs225241970 located at 71,031,384 bp, was recently identified as a *de novo* deletion in B6J that significantly reduced gene expression^38^. When repaired using CRISPR-Cas9 editing, it fully restored expression of *Gabra2* in B6J mice^38^. This finding is consistent with our observation on *Gabra2* expression and sequence variation. We confirmed the presence of this indel variant rs225241970 in the B6J mice used for the RNA sequencing study, and observed that B6J is the only founder strain with a deletion in the 8 founder strains in the CC (Fig. 6c). We also queried this region using the msBWT tool, and determined that the variant does not create alternative splicing of the transcript. Overall, this suggests that the intronic variant in B6J is private to that founder strain and causes a reduction in expression of *Gabra2*, which leads to enhanced seizure sensitivity consistent with the allelic effect for *Ccmss2*/*Ccgss2* identified in QTL mapping (Table. 1).

## DISCUSSION

We present new mouse model resources to guide the identification of genetic variants conferring risk/susceptibility for multiple phenotypic frontiers of epilepsy. We identified novel epileptic mouse strains with extreme epileptic responses that include but are not limited to: 1) multiple CC strains with extreme seizure susceptibility; 2) two CC strains that exhibit resistance to seizure propagation; 3) four CC strains that exhibit SUDEP, possibly with independent mechanisms; 4) CC strains with resistance to epileptogenesis as measured by kindling. Together, these CC strains provide novel animal models to explore molecular, cellular, and physiological basis of various manifestations of epilepsy. We identified a larger phenotypic range and more reliable measures of seizure susceptibility and kindling compared to the Hybrid Mouse Diversity Panel^39^, showing that the CC population harbors a large number of unstudied genetic variants associated with seizure sensitivity and development. The identification of new mouse models with extreme epileptic response makes the CC well suited to serve as a model for the development and deployment of targeted therapeutics for precision medicine in epilepsy^40^. We also have the ability to create congenic mouse strains with distinct sensitivity loci and test targeted therapeutic options that are designed for a particular genetic variant associated with epilepsy. This also allows us to test how the genetic background interacts with a candidate variant and predicted pharmacological interventions.

Among the extreme seizure responses we identified, SUDEP is the fatal complication of epileptic individuals (∼1:1,000/year). There are no current methods that effectively predict or prevent SUDEP^3^. The rare and unpredictable nature of SUDEP restricts traditional GWAS in humans. Respective reviews using postmortem reports have identified some candidate genes, but these studies suffer from a low number of cases and unclear causes of death^7^. The limited genetic and molecular mechanistic insights, coupled with the near absence of suitable animal models of SUDEP, have hindered the development of preventions. Here we have identified four CC strains (CC003, CC008, CC009, and CC029) exhibiting high risk of death following a single episode of transient seizure otherwise nonfatal to classical inbred mouse strains. The finding that younger CC009 mice were more susceptible to seizure-induced sudden death compared to older mice is consistent with clinical observations that early age is an epidemiological risk factor of SUDEP^32,41.^ The causes of age sensitivity in SUDEP are not yet known, but require further study.

Seizure propagation is the process by which a focal seizure spreads within the brain and cause more severe behavioral manifestations. All strains we tested showed progression of seizures from transient involuntary jerks (i.e. MS) to full-blown continuous generalized clonus (i.e. GS) after flurothyl exposure^42^. The decoupling of MS and GS in CC058 and CC032 suggests the genes segregating in these strains can control the spread of seizure activity into contiguous areas via local connections. Identification of distinct genes responsible for generalized seizure propagation may provide pharmacological targets, thereby reducing the harmful and life-threatening complications related to more severe forms of generalized seizures. Besides seizure propagation, seizure development (i.e. epileptogenesis) is another critical yet less understood domain of epilepsy. Epileptogenesis can be modeled in rodents by kindling, a process whereby repeated seizures or subconvulsive stimuli lead to increases in severity and/or susceptibility to subsequent seizures^43,44.^ The kindling process indicates a long-lasting plasticity of the neuronal excitability mimicking epilepsy development^45^. We found lack of kindling effects in many CC strains, including those with intermediate initial seizure susceptibility, such as CC055 and CC051, suggesting resistance to epileptogenesis is not a consequence of a floor effect of the flurothyl kindling model (Supplemental Fig. 4). The identification of the mouse models that exhibit resistance to epileptogenesis allows QTL mapping of F2 progenies of kindling resistant and susceptible strains, thus providing insights into the genetic mechanism of epileptogenesis and revealing druggable targets for preventive therapies.

By combining a high throughput seizure induction model with a mapping population derived from the CC genetic reference mouse population, we were able to identify several genetic associations with seizure sensitivity. As a proof-of-concept, our approach validated an intronic variant exclusively found in the B6J substrain in *Gabra2* as a strong candidate for the transgressive QTL on chromosome 5, notwithstanding that flurothyl screening may have biased sensitivity to phenotypes regulated by GABAR-dependent mechanisms. While here we limit our discovery to a proof-of-concept, the resources we provide here should enable the discovery of genes linked to complex seizure traits, including those such as resistance to epileptogenesis which have, before now, lacked appropriate animal models. Importantly, the resources and tools are publically available and supported by the Systems Genetics Core Facility at UNC (see Data Availability), making the identification of candidate risk/susceptibility genes easily accessible to the scientific community through the targeted generation of F2 populations bred from strains with phenotypic extremes.

## Methods

See Supplemental Methods for experimental details.

## Acknowledgements

This work was supported in part by the following grants from the National Institutes of Health: 4U19AI100625 (FPMV), 1P01AI132130 (FPMV), 5U42OD010924 (FPMV), U24HG10100 (FPMV), R01HD093771 (BDP), T32HD040127 (JRS), and the Rett Syndrome Research Trust (LHW, FPMV). The Systems Genetics Core Facility (provider of CC mice) is partially supported by the University Cancer Research Fund granted to Lineberger Comprehensive Cancer Center (MCR012CCRI). We would like to thank Dr. Marty Ferris, Dr. Wesley Burks, and Dr. Michael Kulis (UNC) for sharing mice in the F2 study as part of the TTSAQ17P1 (FPMV).

## Author contributions

B.G., J.R.S., L.H.W., B.D.P and F.P.V. designed the experiment, discussed and interpreted the findings, and drafted the manuscript. B.G. and L.H.W. performed the *in vivo* experiments. J.R.S performed the QTL mapping. J.R.S and B.G. performed bioinformatics analysis. T.A.B. and P.H. performed DNA extraction. B.G., L.H.W., K.A.D and Y.P. reviewed and analyzed the *in vivo* data. D.R.M. and G.D.S. managed and coordinated the distribution of the CC strains. B.C.C. performed ECG and echocardiography. B.D.P. and F.P.V. supervised the study.

