## Supplemental Material for "Collaborative Cross Mouse Populations as a Resource for the Study of Epilepsy"

### METHODS

#### Mice

Mice (postnatal day 50–90; P50–P90) were studied across four cohorts. Cohorts 1 and 4 represent mice used for screening multiple flurothyl-induced seizure phenotypes across 35 CC strains and 2 classical inbred strains. Cohort 1 consisted of the following strains: CC001/Unc, CC003/Unc, CC004/TauUnc, CC008/GeniUnc, CC009/Unc, CC019/TauUnc, CC021/Unc, CC022/GeniUnc, CC025/GeniUnc, CC027/GeniUnc, CC037/TauUnc, CC038/GeniUnc, CC052/GeniUnc, CC053/Unc, CC055/TauUnc, CC058/Unc, CC071/TauUnc, CC075/Unc, CC076/Unc, C57BL/6J (B6J), and DBA/2J. Cohort 2 represents mice used for the genetic mapping study and includes strains CC027, B6J, their F1s and F2s born in 2016. Cohort 3 represents separate cohort of CC027 and B6J used for the RNA sequencing study in 2018. Cohort 4, screened after completion of cohort 1, consisted of an additional 16 CC strains: CC012/GeniUnc, CC013/GeniUnc, CC024/GeniUnc, CC029/Unc, CC031/GeniUnc, CC032/GeniUnc, CC036/Unc, CC039/Unc, CC040/TauUnc, CC041/TauUnc, CC044/Unc, CC046/Unc, CC051/TauUnc, CC078/TauUnc, CC080/TauUnc, CC081/Unc. Hereafter, CC strains are abbreviated by their strain number (CC001). All CC mice were acquired from the Systems Genetics Core Facility at the University of North Carolina (UNC) at Chapel Hill[1] and were born between 2016–2018. B6J mice used for cohorts 1 and 2 were obtained from a colony maintained by the Pardo-Manuel de Villena laboratory for five generations, and were originally purchased from the Jackson Laboratory (Bar Harbor, ME) in 2016. DBA/2J mice used in cohort 1 were purchased directly from Jackson Laboratory in 2016. Cohort 3 B6J mice used for the RNA sequencing experiment were purchased from Jackson Laboratory in 2018. All mice were raised on standard mouse chow and kept on a 12:12 light/dark cycle. All mouse work was compliant with UNC Institutional Animal Care and Use Committee protocols.

#### Flurothyl-induced seizure and flurothyl kindling

Each mouse was placed in a 2 liter glass chamber inside of a chemical fume hood and allowed to habituate for 1 min before the chamber was closed and 10% flurothyl (bis-2,2,2-trifluoroethyl ether; Sigma-Aldrich) in 95% ethanol was infused at a rate of 200 µL/min onto filter paper (Whatman, Grade 1) suspended at the top of the chamber[2]. Mice exhibit various stages of increasing seizure severity in response to flurothyl exposure, including myoclonic seizure (sudden involuntary jerk/shock-like movements involving the face, trunk, and/or limbs) and generalized seizure (also known as clonic-forebrain seizures that are characterized by clonus of the face and limbs, loss of postural control, rearing, and falling). Generalized seizures can immediately progress into brain stem seizures manifested by tonic extension of the limbs[3]. Upon emergence of a generalized seizure, the lid of the chamber was immediately removed, allowing for rapid dissipation of the flurothyl vapors and exposure to fresh air. Mice were returned to their home cage following recovery from seizures. One mouse at a time was tested in the flurothyl chamber, which was recharged with fresh filter paper, cleaned using water, and thoroughly dried between subjects. B6J mice were challenged in each round of experiments and exhibited similar seizure susceptibility, therefore ensuring lack of differences that could arise from, for example, different batches of flurothyl or different experimenter and video reviewer.

For flurothyl kindling, flurothyl exposures were repeated once daily over eight consecutive days. Mouse behavior during each flurothyl exposure was video-recorded and reviewed by investigators (blind to strains) who determined latency to the onset of both myoclonic and generalized seizures. Mice that failed to survive through the eight-day kindling were excluded from analysis. The linear regression of day-1 through day-8 was used to estimate the kindling effect as previously described[4,5].

**Pentylenetetrazol (PTZ) induced acute seizure and PTZ kindling**

For PTZ-induced acute seizure, each mouse was subjected to PTZ injection (40 mg/kg, i.p.) and animal behavior was monitored by video for 30 min. Video was later analyzed by investigators, blinded to strain, who measured the latency to the onset of behavioral seizures (seizure threshold) and determined seizure severity using a modified Racine's scale: Class 0 = no response, resting; Class 1 = ear and facial twitching, rigid posture with extended tail; Class 2 = transient twitching axially through the body; Class 3 = forelimb clonus and/or rearing; Class 4 = loss of posture control, rearing and falling; Class 5 = jumping and running, generalized tonic-clonic seizure; and Class 6 = death. If the mouse did not exhibit any behavioral seizure within 30 min after PTZ injection, latency was recorded as 1800 s. The maximum behavioral seizure score was measured every 2 min, and the cumulative seizure score was summed. For PTZ kindling, mice received 35 mg/kg PTZ (i.p.) 10 times delivered every other day (on $1^{st}$, $3^{rd}$, $5^{th}$, $7^{th}$, $9^{th}$, $11^{th}$, $15^{th}$, $17^{th}$, $19^{th}$, and $21^{st}$ days of the study). The maximum behavioral seizure score within 30 min after each PTZ injections was assessed using modified Racine's scale described above.

**Repeated-low dose kainic acid (KA) induced seizures**

A repeated-low dose KA model of epilepsy was adopted in this study to evaluate seizure susceptibility involving the limbic system. KA (Sigma-Aldrich) was prepared fresh in sterile distilled water at a concentration of 2 mg/ml, and injected (5 mg/kg, i.p.) once every 30 min until the onset of Class 5 seizures characterized by generalized tonic-clonic convulsions with lateral recumbence or jumping and wild running followed by generalized convulsions. Latency to the onset of Class 5 seizures was determined by investigators who were blinded to mouse strain.

**Electrocardiography (ECG) and echocardiography (Echo) monitoring in mouse**

ECG and Echo recordings were performed by Dr. Brian C. Cooley in the Animal Surgery Core Lab of the McAllister Heart Institute at UNC. Echo was measured on a gently restrained conscious mouse using a Vevo 2100 ultrasound (FUJIFILM VisualSonics Inc.). Parasternal views were used to obtain M-mode images for measuring (1) left ventricle wall thicknesses, (2) internal diameters at end-systole and end-diastole, and (3) ejection fraction and fractional shortening. A representative film of the ultrasound was selected and 6 heartbeat cycles of each measurement were taken to get an average per mouse. ECG was done with an Indus rodent system on mice anesthetized with isoflurane. We used 3-point contact for obtaining electrical signals of the cardiac rhythm and for determining heart rate, contraction, and relaxation times. A representative heart pattern was selected and approximately 1000 to 1200 sequential heartbeats were processed and averaged. QT interval was measured from the beginning of the Q wave until the T wave returned to the isoelectric baseline. Since the QT interval covaries with the RR interval, we calculated a rate-corrected QT interval ($QT_c$) using the formula: $QT_c = QT/(RR/100)^{½}$ [6].

**Genotyping**

Tail biopsies were taken from mice following euthanasia, and DNA was extracted using the QIAGEN DNeasy kit as per manufacturer's instructions. MiniMUGA by GeenSeek (Neogen, Lansing, MI) was used to genotype all mice used in this study in 2018 (https://genomics.neogen.com/en/mouse-universal-genotyping-array). The MiniMUGA array, based on the Illumina Infinium platform, contains 9,914 (prior to 2019) or 10,880 (2019 or thereafter) SNP markers. Informative markers were selected by taking markers that discriminated between B6J and CC027. Next, we identified the frequency of these markers in the F2 population and tested for expected Mendelian ratios. Markers that significantly deviated from the expected frequency distribution were removed from the initial quantitative trait loci (QTL) mapping analysis. The final list of SNP markers from MiniMUGA used for analysis was pruned to 2,440.

After initial analysis, we identified an additional SNP marker on chromosome 5 at 78,300,721 bp to discriminate between the B6J haplotype in CC027 and the B6J from the other parental strain. This variant was discovered using WGS from the CC msBWT tool (http://www.csbio.unc.edu/CEGSseq/index.py)[7-9]. PCR primers were designed to amplify a 431bp DNA fragment containing the SNP of interest: Forward–CTGATGTCCAGATTGCTTAGT, position 78,300,580; Reverse–GGATTTGAGAAGGAAGCTAGAA, position 78,301,010. PCR was performed under the following conditions: 95°C for 2 min, 35 cycles of 95°C, 55°C, 72°C for 30 sec each, followed by a 72°C hold for 7 min. PCR reactions were visualized on 2% agarose gels with ethidium bromide. Each PCR reaction contained 1 µl crude genomic DNA, 2 µl 5X PCR buffer (Promega, Madison, WI), 1 µl dNTP mix (2.5 mM each), 0.3 µl of each primer, 0.1 µl Taq (GoTaq DNA polymerase, Promega) and $H_2O$ for a total of 10-µl reaction volume. PCR products were genotyped by sequencing at Eurofins Genomics. 140 F2 samples that had ambiguous parental origin were genotyped to resolve the QTL boundary on chromosome 5.

**Heritability estimation**

Heritability is the proportion of the overall variability observed in a trait that is due to inherited genetic factors. We calculated a marker-based estimation of heritability that included a kinship relatedness matrix based on genotypes using the R package heritability[10]. The kinship matrix was created using genotype probability files of 35 CC strains from genome build 38 (http://csbio.unc.edu/CCstatus/index.py?run=FounderProbs) with Rqtl2[11].

**QTL mapping**

We performed QTL mapping using the R packages, Rqtl and Rqtl2[11,12]. Rqtl2 performs QTL mapping through a regression of the phenotype at each marker. Rqtl was used to perform multiple-QTL mapping (MQM), which tests for additive and interacting QTL effects. Age was included as a covariate for trait mapping. QTL significance intervals were defined by the 95% Bayesian credible interval, calculated by normalizing the area under the QTL curve[13]. Log of the odds ratio (LOD) was the reported mapping statistic. The significance thresholds for QTL were calculated using 10,000 permutations.

**Tissue preparation**

Fourteen mice were anesthetized with euthasol (100 mg/kg, i.p.) prior to decapitation and brain removal. We dissected the hippocampus, cortex, and cerebellum in addition to lungs, livers, and tails from eight B6J (four males, four females) and six CC027 (four males, two females) that were between 80–100 days old. Tissue was flash frozen on liquid nitrogen and stored at -80$^{o}$C before RNA extraction.

**RNA sequencing**

RNA sequencing was performed on hippocampus tissue for fourteen mice. RNA was submitted to the UNC High Throughput Sequence Facility for library preparation and sequencing. All samples had a RIN score > 8.0 and an initial concentration > 90 ng/ul. NuGEN Universal Plus mRNA-Seq was used for library preparation. Sequencing was performed on the Illumina HiSeq 4000 with 50 bp stranded single end reads and pooled libraries across 2 lanes at an average sequencing depth of 9 million reads per sample. The short read aligner BBmap was used to align reads to the MM10 mouse reference genome. DEseq2 was used to quantify transcript expression levels and test for differential expression[14]. “Genotype” was used as the main term for the DEseq2 generalized linear model.

RNA sequence reads were also converted into msBWTs and are available at http://www.csbio.unc.edu/CEGSseq/index.py. This tool was used to identify and validate sequence differences of transcripts between CC027 and B6J.

**Characterization of QTL by homologous genes in humans**

Human orthologous genes within each QTL interval were identified using BioMart query in Mouse genes (GRCm38.p6) dataset. Per QTL, we compared the human orthologous genes with those reported in the EpilepsyGene database[15].

**Data Availability**

Genotypes for MiniMUGA are available at https://www.med.unc.edu/mmrrc/genotypes. CC sequenced genomes can be queried using the msBWT tools available at http://www.csbio.unc.edu/CEGSseq/index.py?run=MsbwtTools. Phenotype data from this study is available on the Mouse Phenome Database at https://phenome.jax.org/projects/Shorter9 with accession number MPD:660. Zenodo.org accession no. 3250238 also provides access to the pruned genotypes from MiniMUGA, FASTQ files used for RNAseq analysis, phenotypes for the CC strains, and phenotypes for the F2 mapping population.

### SUPPLEMENTAL TABLES AND FIGURES

| QTL | Chr | Gene symbol | Description | Phenotypes |
|---|---|---|---|---|
| *Ccmss1* | 4 | *NKAIN3* | Na+/K+ transporting ATPase interacting 3 | SMEI |
| | | *COQ3* | coenzyme Q3 methyltransferase | EE |
| *Ccmss2/Ccgss2* | 5 | *GABRB1* | gamma-aminobutyric acid (GABA) A receptor, beta 1 | EE |
| *Ccmss3/Ccgss3* | 10 | *PTPRR* | protein tyrosine phosphatase, receptor type, R | EE |
| *Ccmss4* | 13 | N/A | N/A | N/A |
| *Ccgss1* | 1 | *SLAMF1* | signaling lymphocytic activation molecule family member 1 | EE |
| | | *ATP1A2* | ATPase, Na+/K+ transporting, alpha 2 polypeptide | FS/GEFS+ |
| | | *IGSF8* | immunoglobulin superfamily, member 8 | SMEI |
| | | *KCNJ10* | potassium inwardly-rectifying channel, subfamily J, member 10 | EAST syndrome |
| *Ccgss4* | 11 | *RAPGEF6* | Rap guanine nucleotide exchange factor (GEF) 6 | EE |

Abbreviation: SMEI = severe myoclonic epilepsy in infancy; EE = epileptic encephalopathy; FS = febrile seizures; GEFS+ = generalized epilepsy with febrile seizures plus.

**Supplemental Table 1. Human homologous genes in QTL overlapped with genes relevant to human epilepsy archived in EpilepsyGene database.**

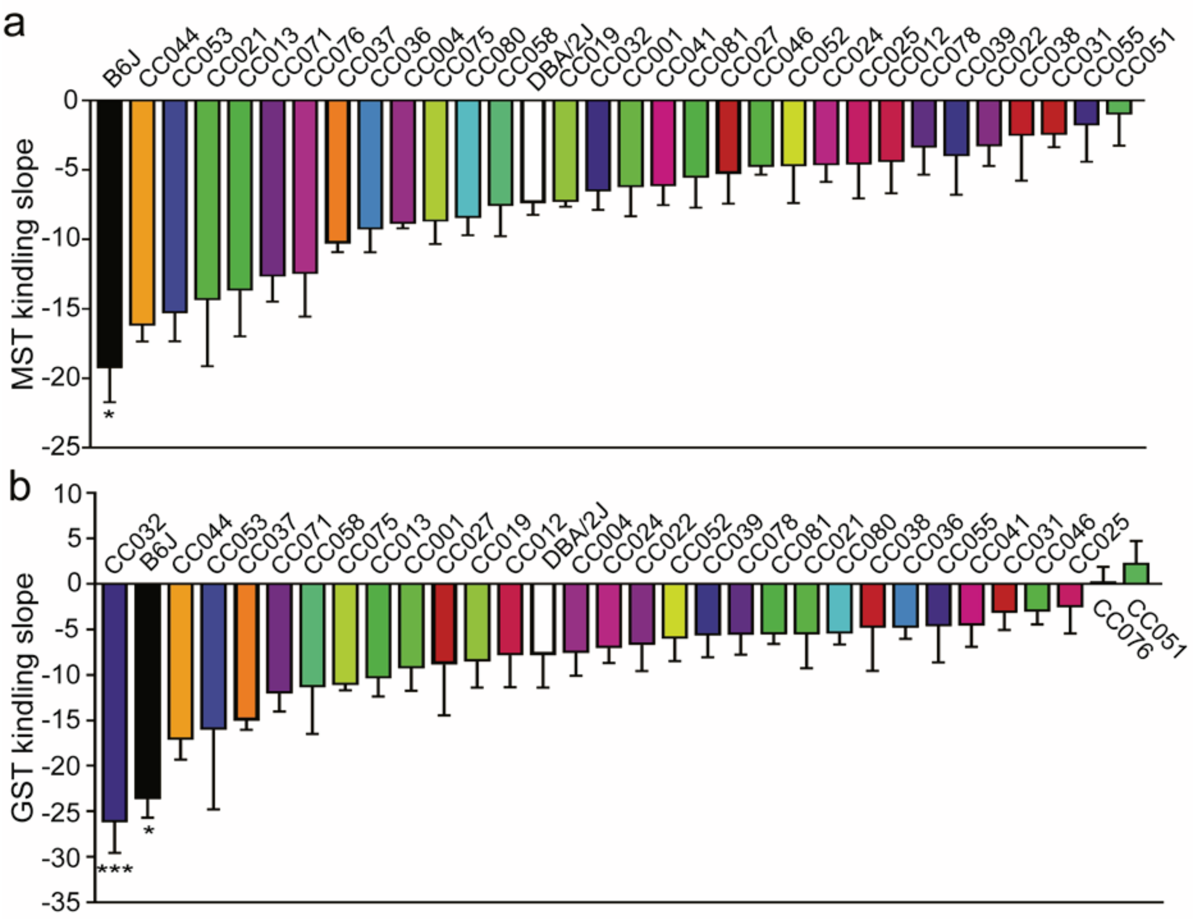


**Supplemental Figure 1. Strain dependence of flurothyl kindling slope. a**, MST and **b**, GST 8-day flurothyl kindling slope. Data are presented as mean ± SEM and analyzed using one-way ANOVA with *post hoc* Dunnett's multiple comparisons test, n=3–10. *p<0.05 and ***p<0.001 compared to DBA/2J.

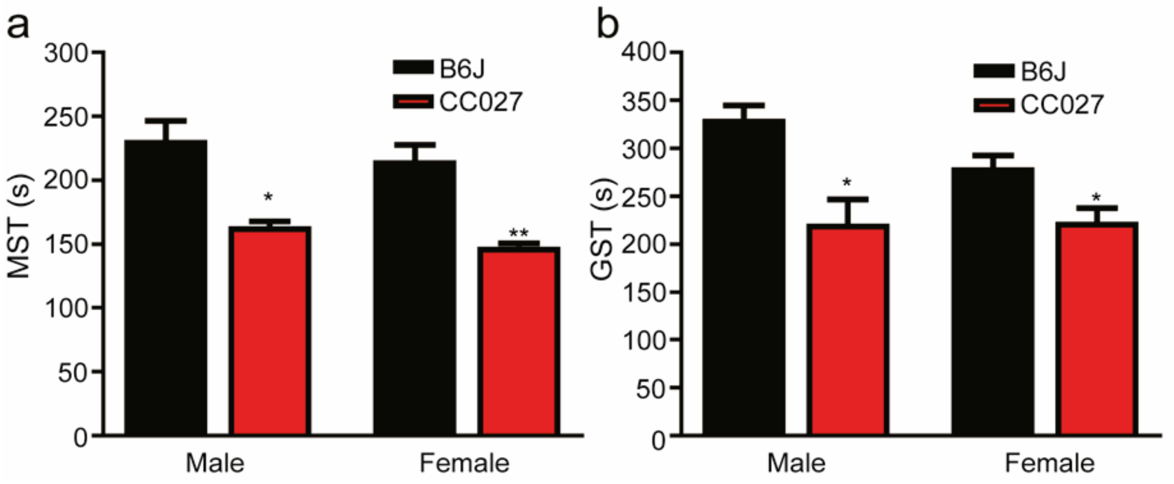


**Supplemental Figure 2. Lack of sex effect on seizure threshold in B6J and CC027 mice.** **a**, MST and **b**, GST measurements. Data are presented as mean ± SEM and analyzed using two-way ANOVA with *post-hoc* Dunnett's multiple comparisons test, n=4. *p<0.05 and **p<0.01 compared to B6J.

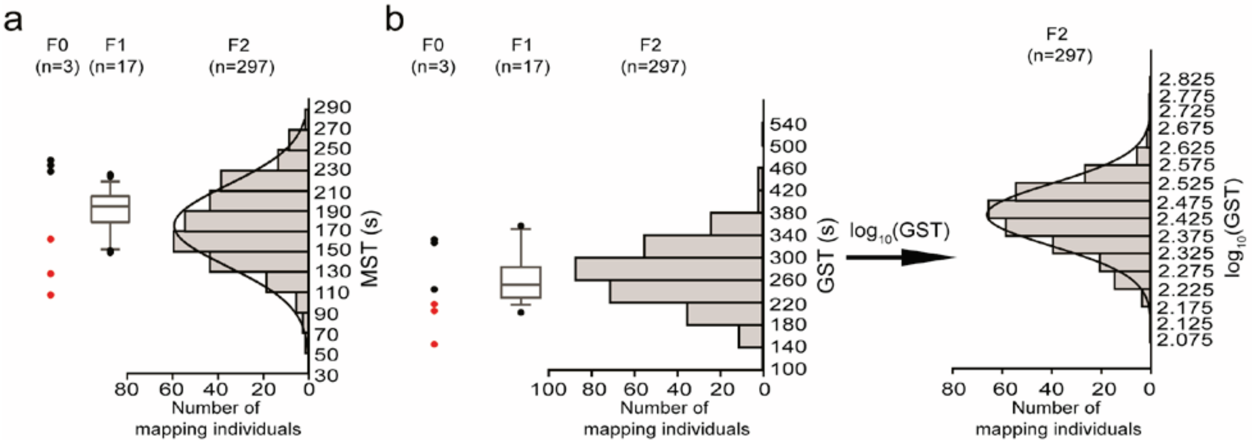


**Supplemental Figure 3. Seizure thresholds in F0, F1, and F2 generations.** Frequency distribution of **a**, MST, **b**, GST and its $\log_{10}$ transformation in F2 mapping population (n=297) derived from the cross between the seizure resistant B6J (F0, black dots, n=3) and seizure susceptible CC027 (F0, red dots, n=3) in parallel with F1 (n=17, median with 10–90 percentile).

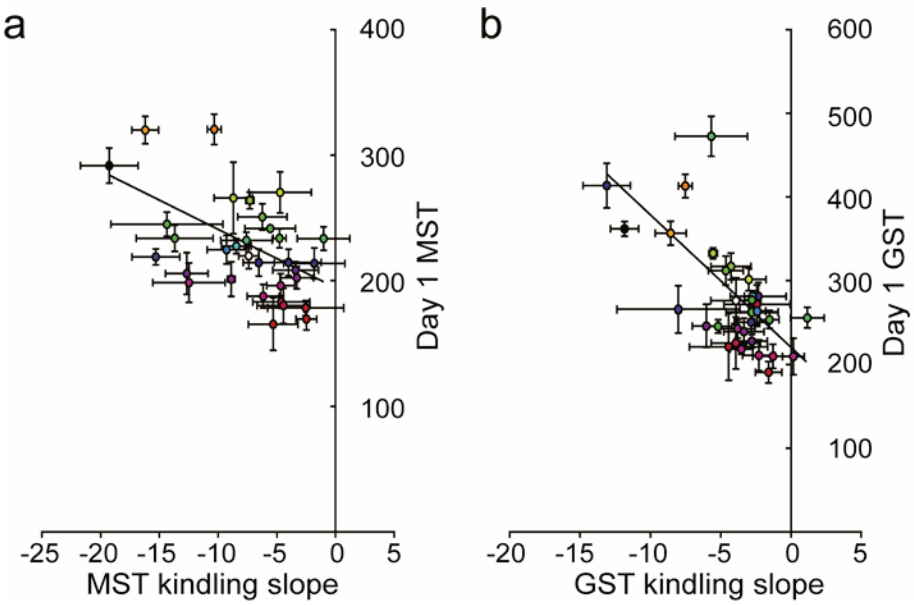


**Supplemental Figure 4. Correlation of kindling slope and day 1 seizure susceptibility.** Correlation of 8-day kindling slope and day 1 seizure threshold in the measurement of **a**, MST ($R^2$=0.263, p<0.01) and **b**, GST ($R^2$=0.442, p<0.001). Data are presented as mean ± SEM.
